# All-Atom MD Simulations of the HBV Capsid Complexed with AT130 Reveal Secondary and Tertiary Structural Changes and Mechanisms of Allostery

**DOI:** 10.1101/2021.02.08.430329

**Authors:** Carolina Pérez Segura, Boon Chong Goh, Jodi A. Hadden-Perilla

**Affiliations:** Department of Chemistry and Biochemistry, University of Delaware, Newark, Delaware 19716; Antimicrobial Resistance Interdisciplinary Research Group, Singapore-Massachusetts Institute of Technology Alliance for Research and Technology Centre, Singapore 138602

**Keywords:** hepatitis B virus, HBV, virus capsid, core protein allosteric modulator, CpAM, phenyl-propenamide, AT130, molecular dynamics simulations, physical virology, computational virology

## Abstract

The hepatitis B virus (HBV) capsid is an attractive drug target, relevant to combating viral hepatitis as a major public health concern. Among small molecules known to interfere with capsid assembly, the phenylpropenamides, including AT130, represent an important anti-viral paradigm based on disrupting the timing of genome encapsulation. Crystallographic studies of AT130-bound complexes have been essential in explaining the effects of the small molecule on HBV capsid structure; however, computational examination reveals that key changes attributed to AT130 were erroneous, likely a consequence of interpreting poor resolution arising from a highly flexible protein. Here, all-atom molecular dynamics simulations of an intact AT130-bound HBV capsid reveal that, rather than damaging spike helicity, AT130 enhances the capsid’s ability to recover it. A new conformational state is identified, which can lead to dramatic opening of the intradimer interface and disruption of communication within the spike tip. A novel salt bridge is also discovered, which can mediate contact between the spike tip and fulcrum even in closed conformations, revealing a mechanism of direct communication across these domains. Combined with dynamical network analysis, results describe a connection between the intra- and interdimer interfaces and enable mapping of allostery traversing the entire capsid protein dimer.

## 1. Introduction

Hepatitis B virus (HBV) is a DNA hepadnavirus around 45 nm in diameter [1]. It causes chronic liver infection in 250 million people worldwide [2] and remains a major public health concern, despite the availability of a vaccine and therapeutic options [3]. The virus is enveloped, and its core is composed of a genome-containing capsid. The capsid, an attractive drug target, is a protein shell that self-assembles from homodimeric core protein (Cp) (Figure 1a). Particles that go on to be infectious incorporate 120 Cp dimers and exhibit T=4 icosahedral geometry. The two quasi-equivalent positions Cp dimers occupy within the symmetry of the capsid are denoted AB and CD, where A chains form the twelve pentamers and B, C, and D chains form the thirty hexamers [4] (Figure 1b). Full-length Cp is composed of 183 amino acid residues (Cp183) and contains two domains: the assembly domain (Cp149) and the C-terminal domain (CTD). While Cp149 controls the capsid assembly process, the CTD is essential for genome packaging, reverse transcription, and intracellular trafficking [1]. Potentially all of these processes could be inhibited by targeting the capsid for antiviral intervention.

**Figure 1.**
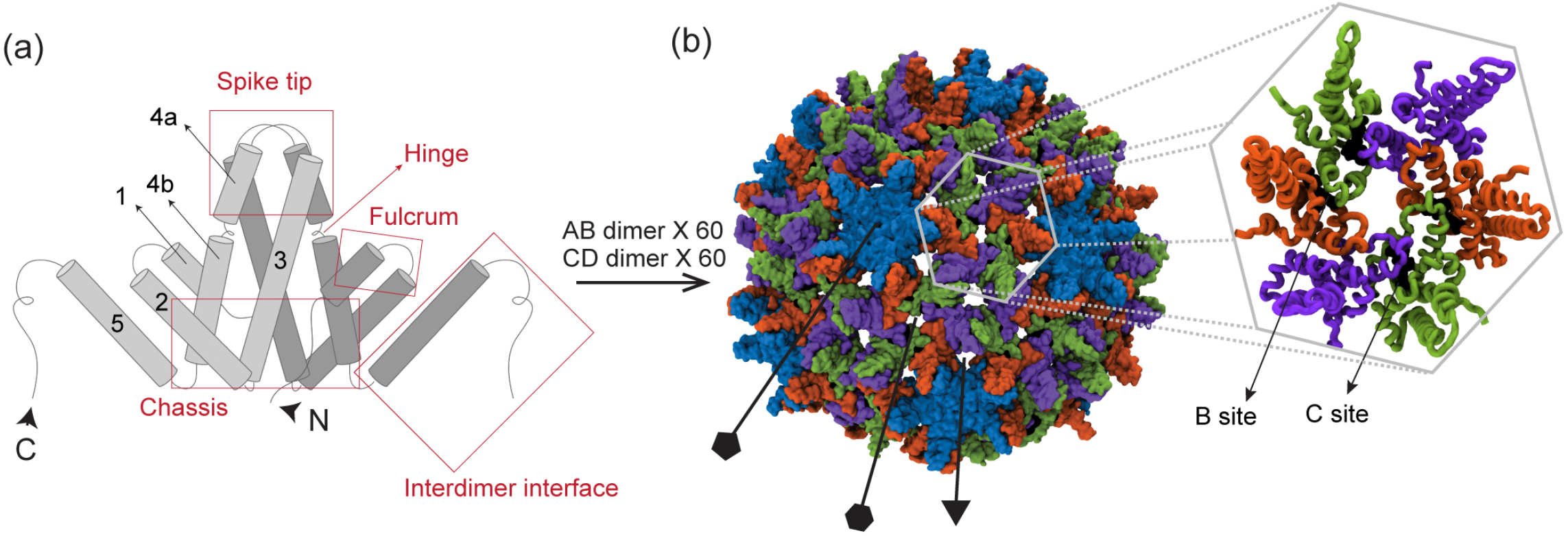
HBV core protein dimer and capsid. (**a**) Schematic of Cp149 dimer, illustrating the five typical *α*-helices, labeled 1-5 from the N-to C-terminus. Established dimer domains [11] are indicated, including the chassis, fulcrum, hinge, spike tip, which forms part of the intradimer interface, and interdimer interface where helix 5 of neighboring chains overlap and form a CpAM binding pocket. (**b**) The T=4 icosahedral capsid incorporates 120 dimers, whose chains can occupy four quasi-equivalent positions, designated A (blue), B (orange), C (green), and D (purple). Capsid-incopororated dimers are thus denoted AB and CD. A chains form the 12 capsid pentamers, while two copies of B, C, and D chains form the 30 capsid hexamers. AT130 binds in B sites, between B and C chains, and C sites, between C and D chains.

Numerous small molecules that affect capsid assembly *in vitro* have been identified. Collectively referred to as Cp allosteric modulators (CpAMs), these small molecules accelerate the assembly process, apparently by binding Cp and modulating its conformation [5]. One such family of CpAMs are the phenylpropenamides (PPAs), which include the compound AT130 [6] (Figure 2a). Unlike CpAMs that also misdirect assembly to produce aberrant structures, the presence of AT130 still leads to the formation of morphologically regular capsids; nevertheless, AT130-bound capsids can become kinetically trapped, fail to correctly enclose the viral genome, and do not become infectious particles [7–9]. Thus, PPAs represent an important anti-viral paradigm based on disrupting the timing of genome encapsidation and nuceolcapsid production, and characterizing their mechanism of action is highly relevant to therapeutic development. AT130 binds the capsid within hydrophobic pockets formed at interdimer interfaces (Figure 1b), particularly those found where the quasi-equivalent C chain overlaps the B chain (B site) and where the D chain overlaps the C chain (C site) [10]. A 4.20-Å crystal structure of the Cp149 capsid saturated with AT130 (120 copies) suggests that changes in the capsid structure are quaternary, with secondary and tertiary disruption in the dimers [10].

**Figure 2.**
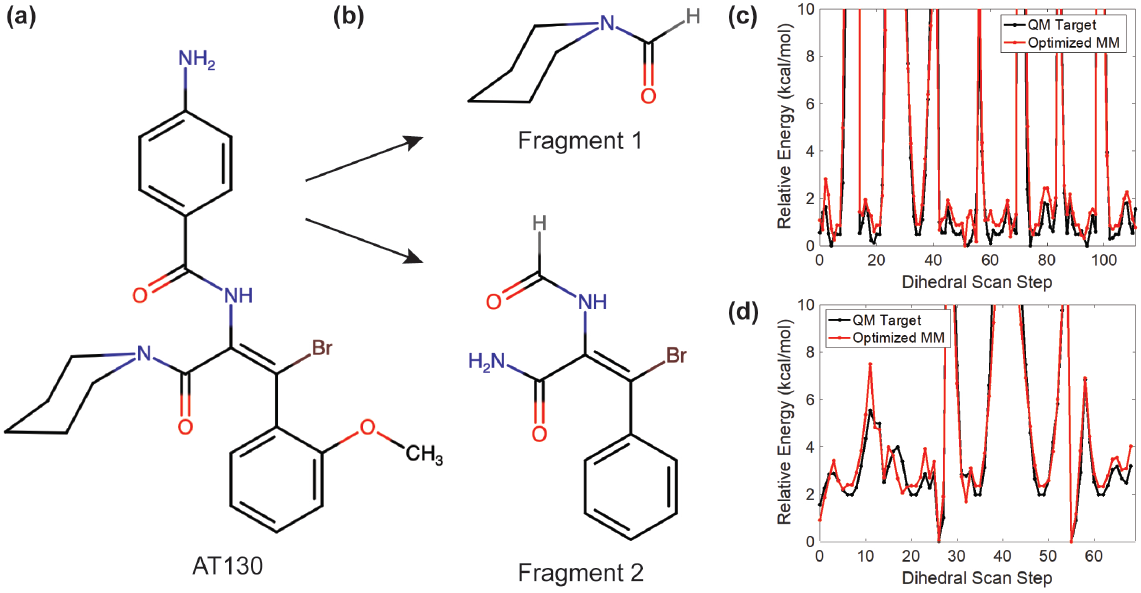
Parameterization of AT130. (**a-b**) The phenylpropenamide derivative AT130 was divided into two fragments for QM calculations. (**c-d**) Optimized dihedral parameters show accurate reproduction of QM rotational profiles.

The extraordinary flexibility of the HBV capsid often renders the atomistic details of its structure challenging to characterize. Owing to the consequences of capsid dynamics, experimental structures tend to exhibit low resolution [12]. All-atom molecular dynamics (MD) simulations offer a robust solution to the limitations of experimental methods and have emerged over the past 15 years as an essential technique for the study of virus structures at a scale applicable to probing their biology [13]. We have previously employed all-atom MD simulations to investigate the intact HBV capsid [12,14,15], as well as the free dimer [16], in all cases revealing deeper and more detailed information on the relationship between structure and function than was accessible to experiments. Additional atomistic studies have investigated early assembly intermediates of Cp149, including in the presence of CpAMs [17–20], providing further insight into the mechanisms of HBV capsid formation and disruption.

Here, we employ all-atom MD simulations to examine an intact AT130-bound capsid and extensively characterize the capsid-incorporated dimer. Our analyses leverage a cumulative 60 *µ*s of sampling each for the AB and CD dimers to overcome the typical convergence challenges associated with computational study of a highly flexible protein. Comparisons are made with an analogous dataset describing the apo-form capsid, extracted from our previous simulation work [12], to definitively demonstrate the effects of AT130 on dimer structure and dynamics under physiological conditions. Our results underscore the power of all-atom MD simulations to elucidate the nature of biological systems in their native environments, providing an indispensable complement to structure determination approaches that focus on static models or single conformations.

Importantly, our findings reveal new information regarding secondary and tertiary response of the dimer to AT130, and indicate that some structural changes previously attributed to the CpAM based on crystallographic data [10] were erroneous. Conclusions based on experimentally-derived models of the AT130-bound complex were likely misled by the poor resolution inevitably obtained for the notoriously flexible capsid. Remarkably, our simulations show that, rather than damaging the helical secondary structure of the spike tip, AT130 enhances the capsid’s ability to rapidly recover it. Consistently folded spike tips lead to smaller average bending angles along the dimer hinge, and, typically, a closed and well-ordered intradimer interface.

However, through extensive sampling, we also identified a conformational state never before observed in experimental structures, characterized by dramatic opening of the intradimer interface and disruption of communication between half-dimers within the spike tip. This rare state was only observed in the CD dimer, occurred more frequently in the quasi-equivalent D chain in the presence of AT130, and often resulted in direct contact of the half-dimer spike tip with the fulcrum via formation of a salt bridge between Arg28 and Asp83. Salt bridge mediated connection of these domains was also found to be possible in dimers whose key internal contacts were maintained, owing to adjustments of helix 2 or 4b, revealing a pathway for allosteric communication between the intra- and interdimer interfaces that does not pass through the four-helix bundle. Both the open spike state and higher populations of above- and below-average dimer hinge angles likely account for the increased conformational variation captured for spikes tips in the AT130-bound crystal model [10], which served to blur resolution and mislead experimental interpretation.

## 2. Materials and Methods

### 2.1. Computational Modeling

An atomistic model of the AT130-bound Cp149 HBV capsid was constructed based on an available crystal structure of the complex, PDB 4G93 [10]. Missing residues at the C-terminus of each protein chain were rebuilt using ROSETTA3 [21]. Residues 143-149 of five A chains forming a capsid pentamer were folded simultaneously to produce 2,000 conformations, or 10,000 total structures for Cp149 in the quasi-equivalent A position. Residues 141-149 of two B chains, residues 140-149 of two C chains, and residues 143-149 of two D chains forming a capsid hexamer were folded simultaneously to produce 5,000 conformations, or 10,000 total structures each for Cp149 in quasi-equivalent B, C, and D positions. The four ensembles were clustered using the partitioning around medoids algorithm based on root-mean-square deviation of the protein backbone [22] for the modeled termini. The medoid of the most populated cluster was selected as the final model for each chain in the capsid asymmetric unit.

The complete, intact capsid structure was generated by applying transformation matricies for a T=4 icosahedron. Hydrogen atoms were added to the model using PDB2PQR [23] for a physiological pH of 7.0, assigning the protonation states of titratable groups according to local pKa. Sodium and chloride ions were placed around the capsid using the cionize module of Visual Molecular Dynamics (VMD) [24]. The capsid, along with local ions, was solvated in a 39.2 nm box of TIP3P water [25] with NaCl concentration of 150 mM. Simulation files were prepared with VMD’s psfgen module, applying the CHARMM36 force field [26].

### 2.2. AT130 Parameterization

Custom force field parameters compatible with CHARMM36 were developed to describe AT130 (Figure 2a). The CHARMM general force field (CGenFF) program [27] was used to assign atom types and partial atomic charges by analogy, as well as to identify missing parameters not covered by CGenFF. The missing bond, angle, and dihedral parameters, those with high penalty scores, and the partial charges of all atoms were optimized using the Force Field Toolkit (ffTK) [28] plugin in VMD and Gaussian09 [29]. The 53-atom AT130 molecule was split into two fragments (Figure 2b) to isolate key parameters and reduce the computational overhead of quantum mechanics (QM) calculations. The final potential energy profiles of refitted dihedral parameters demonstrated agreement with the those computed with QM (Figure 2c-d). All missing charges and parameters optimized using standard ffTK protocols were recombined with CGenFF to yield a complete parameter set for AT130. Notably, CGenFF parameters optimized using ffTK have been independently shown to appropriately model Cp-AT130 interactions and binding stability during MD simulations [30].

### 2.3. Molecular Dynamics Simulations

MD simulations were performed with NAMD 2.13 [31]. The capsid system was subjected to energy minimization with the steepest descent algorithm for 30,000 cycles, then thermalized by increasing the temperature from 60 K to 310 K over a duration of 5 ns. Cartesian restraints of 5 kcal/mol that were previously maintained on the protein backbone were removed over a duration of 5 ns. The capsid system was allowed to equilibrate for 5 ns without restraints prior to collection of 1.0 *µ*s of production sampling. Trajectory frames were written every 20 ps.

MD simulations were performed in the NPT ensemble with a timestep of 2.0 fs. The r-RESPA integrator was used to propagate dynamics. Electrostatic interactions were split between short and long range at a cutoff of 12 Å based on a quintic polynomial splitting function. Long range electrostatics were calculated using the particle mesh Ewald algorithm with a grid spacing of 2.1 Å and 8^*th*^ order interpolation. Full electrostatic evaluations were carried out every other timestep. The Langevin thermostat algorithm was employed for constant temperature control at 310 K with a damping coefficient of 1.0 ps^*−*1^. The Nosé-Hoover Langevin piston algorithm was employed for constant pressure control at 1.0 bar using isotropic scaling with piston oscillation period of 2000 fs and damping timescale of 1000 fs. Bonds to hydrogen were constrained using the SHAKE algorithm for solute and the SETTLE algorithm for water.

### 2.4. Trajectory Analysis

Analysis of MD simulation trajectories was carried out with VMD [24]. Characterization of each dimer system was based on an ensemble of three million total conformations. The 1.0-*µ*s AT130-bound capsid trajectory was deconstructed into sixty 1.0-*µ*s dimer trajectories for AB and CD quasi-equivalent subunits, totalling 60 *µ*s of cumulative sampling for each type of capsid-incorporated dimer. Analogous data for the apo-form capsid was taken from a previously published simulation [12] based on PDB 2G33 [32]. A summary of the simulation systems analyzed in this study is provided in Table 1. The dimer trajectories were aligned to their respective crystal structures using the C*α* atoms of the Cp149 chassis (residues 1 to 10, 26 to 62, and 95 to 110). Secondary structure assignments were made using the STRIDE algorithm [33]. Principal component analysis was carried out as previously described [34], and mode visualization used NMWiz [35]. Twenty modes were calculated based on the capsid’s C*α* trace. Network models were generated in triplicate using three sets of one million randomly selected conformers, with one node per residue mapped to the C*α* atoms. Dynamical network analysis was performed using NetworkView [36]. Curvature of the dimer hinge was measured with Bendix [37], fitting bendices through residues 88-98.

**Table 1.** The four MD simulation systems analyzed in this study.

| apo-form capsid | AT130-bound capsid |
| --- | --- |
| AB capsid-incorporated dimer | AB capsid-incorporated dimer |
| CD capsid-incorporated dimer | CD capsid-incorporated dimer |

## 3. Results

### 3.1. Secondary Structure Analysis

Figure 3 shows a summary of secondary structure adopted by quasi-equivalent A, B, C, and D chains of capsid-incorporated dimers in the apo-form and AT130-bound systems during MD simulations. Notably, the spikes of the apo-form AB dimer are well-folded in the crystal model (PDB 2G33, 3.96 Å [32]), while those of the CD dimer are not (Figure 3, left). Our previous work on the apo-form capsid highlighted the flexibility of the spikes, particularly those of the CD dimer, and increased mobility was shown to correlate with higher B-factors and lower local resolution in experimental data [12]. Nevertheless, disordered spike conformations are in contrast with high resolution structures of the apo-form capsid (PDB 1QGT, 3.30 Å [4] and 6HTX, 2.66 Å [38]), which demonstrate that fully-folded CD spikes are possible.

**Figure 3.**
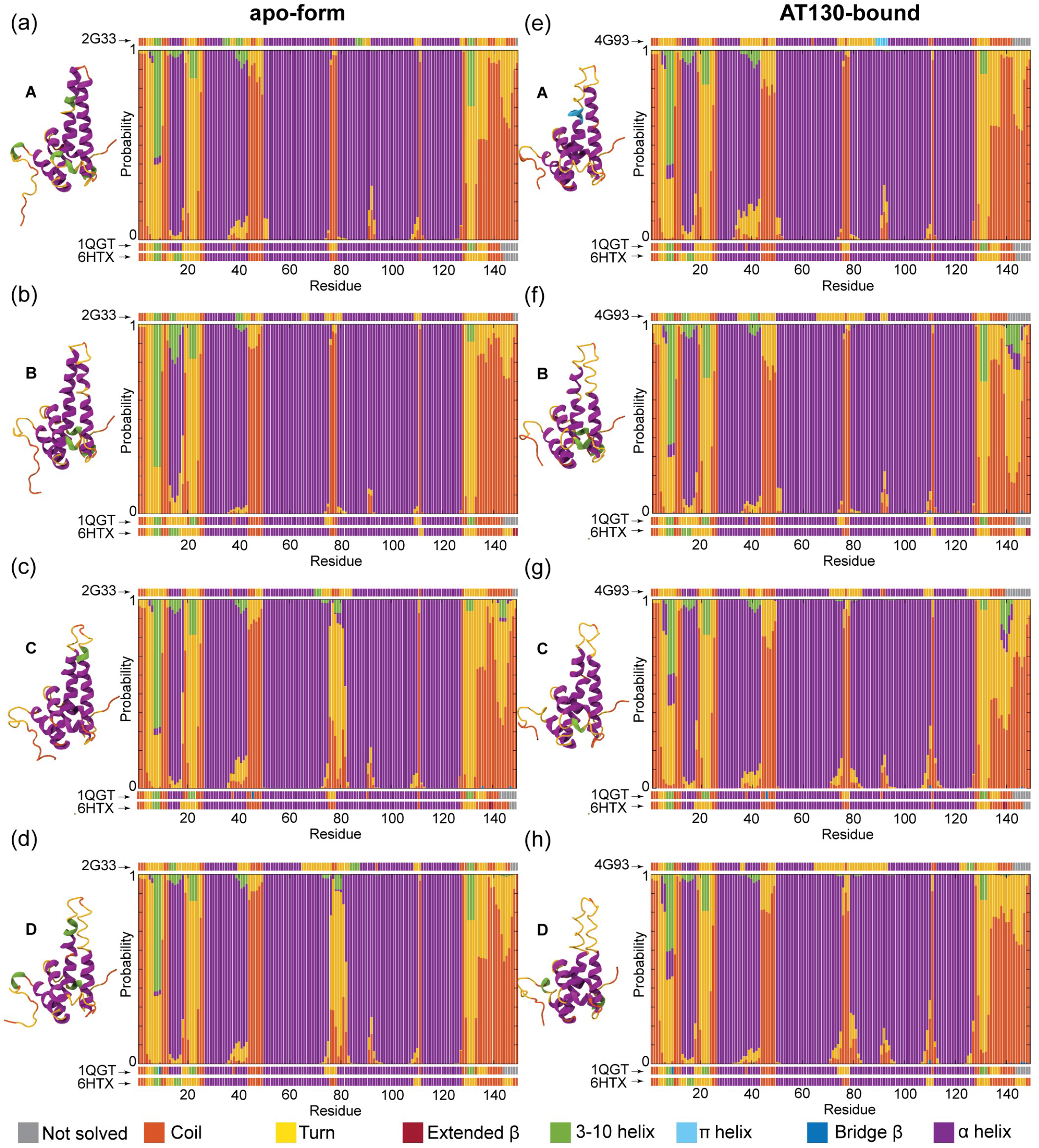
Secondary structure of capsid-incorporated dimers. (**a-d**) Chains A, B, C, and D of apo-form capsid (PDB 2G33, 3.96 Å). (**e-h**) Chains A, B, C, and D of AT130-bound capsid (PDB 4G93, 4.20 Å). Cartoon representation for each chain of crystal models shown, along with their secondary structure assignments according to STRIDE [33]. Histograms indicate probability of each Cp149 residue to be observed in a given secondary structure during MD simulations, calculated based on ensembles of three million conformers collected over 60 *µ*s of cumulative sampling. Purple denotes *α*-helix. Secondary structure of high resolution crystal structures (PDB 1QGT, 3.30 Å [4] and 6HTX, 2.66 Å [38]) provided for reference.

During MD simulations of the apo-form capsid, the spikes of AB dimers were consistently ordered, with residues identified as helical in experimental structures exhibiting a high probability to remain so over microsecond timescales (Figure 3a-b). Taken together with previous results characterizing flexibility [12], these data indicate that spikes can be both well-folded and highly mobile, such that low resolution in experimental structures need not be interpreted as structural disruption. The spikes of CD dimers recovered some helicity relative to the crystal model, particularly in helix 3b, but were far more likely to be observed in partially-folded states (Figure 3c-d and S8a-b). Specifically, a fully-folded spike tip was present in 11.5% and 8.2% of conformers in apo-form C and D chains, respectively, while a spike tip that was folded along helix 3b but partially-folded along helix 4a was present in 75.2% and 79.8% of conformers. Limited simulation timescales entail the consequence that biomolecules retain bias from their initial configurations; given extended sampling, all constituent CD spikes may have eventually refolded. In previous MD simulations of the free dimer, we observed that a salt bridge between Glu77-Arg82 could stabilize partially-folded states of the spike [16]; the salt bridge was identified in 22.7% and 27.5% of conformers in apo-form C and D chains, respectively, accounting in part for their reduced ability to refold.

In the crystal model of the AT130-bound capsid (PDB 4G93, 4.20 Å [10]), the spikes of all four quasi-equivalent chains are disordered, with CD dimers exhibiting even less helical content than in apo-form (Figure 3, right). This effect was attributed directly to the presence of the bound CpAM in the crystallographic study [10]. However, during MD simulations of the AT130-bound capsid, the spikes of AB dimers rapidly recovered helicity, exhibiting nearly the same probability to be fully-folded as in the apo-form capsid (Figure 3e-f). Interestingly, the majority of CD dimer spikes also refolded (Figure 3g-h and S8c-d), with 77.7% and 64.0% of conformers in AT130-bound C and D chains, respectively, found to contain maximum helicity. Ultimately, following equilibration under physiological conditions, the AT130-bound capsid relaxes from the crystal model and adopts significantly more ordered spikes, even in the presence of the CpAM. This finding indicates that AT130 does not damage the helical nature of capsid spikes, but actually enhances the ability of spikes to refold relative to apo-form.

Beyond behavior of the capsid spikes, other differences between the crystal models and MD simulation ensembles are notable (Figure 3). The 310-helical element preceding helix 1, which was not consistently identified in crystal structures, was observed in around half of the ensembles; this element demonstrated a higher probability to exist as an *α*-helix in the AT130-bound capsid. Helix 1, which is not clearly defined in the A, B, and D chains of the highest resolution crystal (PDB 6HTX, 2.66 Å [38]), was shown to exist as the expected *α*-helix the majority of the time in both apo-form and AT130-bound systems. Although the degree of disorder at the C-terminus of helix 2 varied, it occurred with the highest probability in chain A of the AT130-bound capsid. While a break in secondary structure associated with the dimer hinge between helix 4a and 4b was not emphasized in crystallographic data, it can be observed in MD simulation results in the 88-98 residue range. The length of the loop separating helices 4 and 5 exhibited some variability in length in the AT130-bound CD dimer, which is contacted by a CpAM on either side. The C-terminal tail exhibited some unexpected probability to contain 310-helical regions, particularly around residue 140 in the B and C chains containing AT130, beyond the point at which the crystal model was well-resolved.

### 3.2. Dynamical Network Analysis

Figure 4 shows dynamical network models for the AB and CD capsid-incorporated dimers in the apo-form and AT130-bound systems, colored by communities and weighted by the extent of correlation in residue motions. Network models were calculated in triplicate, splitting each ensemble of three million frames into three sets of one million randomly selected frames. Notably, the three sub-datasets produced similar, yet inequivalent results (Figure S9). The observation that 20 *µ*s of conformational sampling was insufficient for precision in the analysis underscores the highly flexible nature of the Cp dimer, and raises the question of convergence in any state-of-the-art computational study that examines shorter timescales. Nevertheless, the presented network models reveal important aspects of correlated motion and communication in the functional dynamics of Cp, and findings are consistent with experimental characterizations of dimer domains based on concerted structural displacements observed in crystal structures [11].

**Figure 4.**
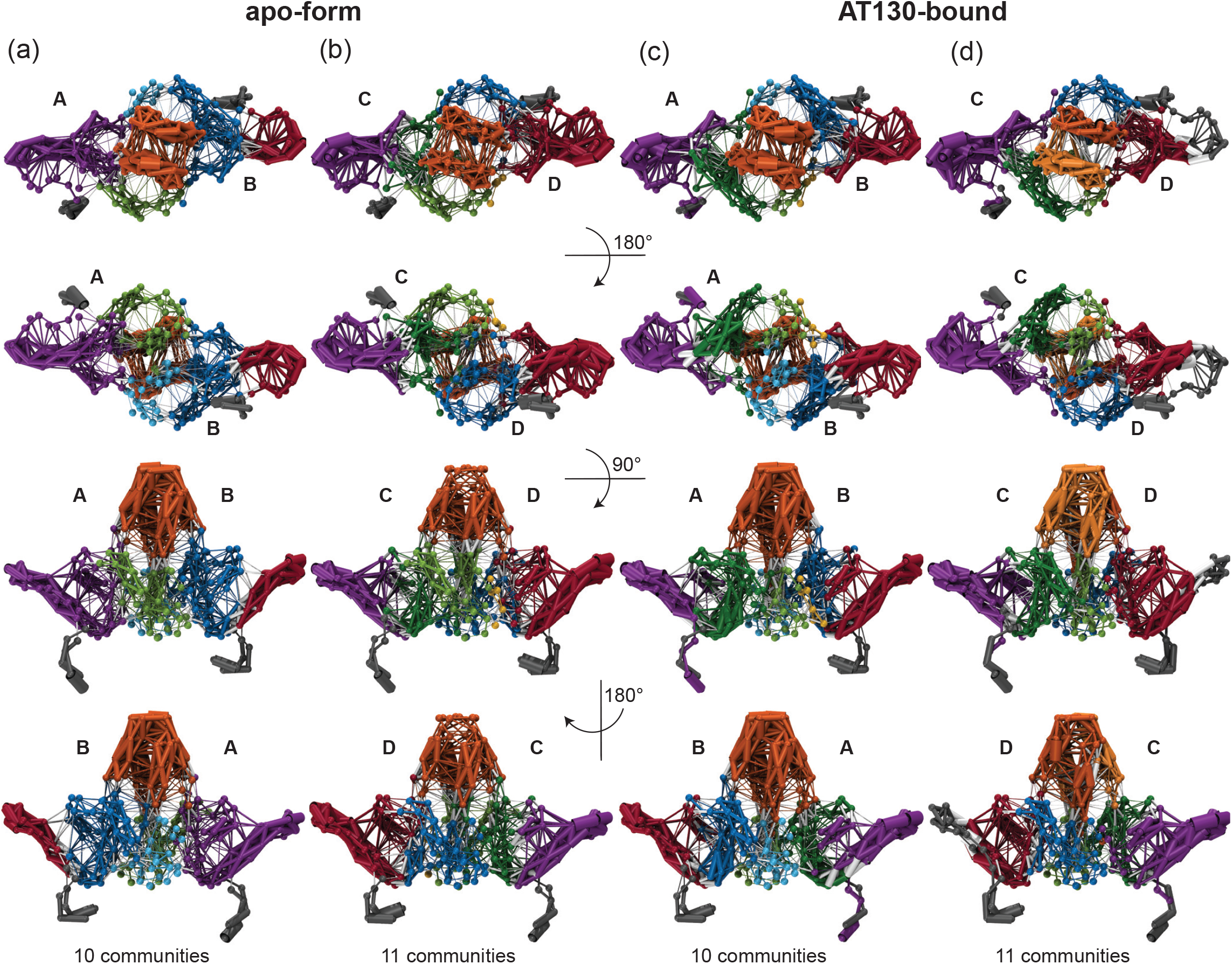
Dynamical network models of capsid-incorporated dimers. (**a-b**) Representative model for apo-form AB and CD dimers. (**c-d**) Representative model for AT130-bound AB and CD dimers. Models calculated in triplicate (Figure S9) based on ensembles of one million conformers collected over 20 *µ*s of cumulative sampling. Models shown colored by communities and weighted by the extent of correlation in residue motions; visualization created with NetworkView [36]. Five major communities encompass helix 1,5 of each half-dimer (purple and red), where helix 5 forms the interdimer interface, helix 2-3a of each half-dimer (blue and green), which are part of the chassis domain, and helix 3b-4a of both half dimers (orange), called the spike tip. The remainder of the fulcrum domain participated in either the helix 1,5 or helix 2-3a community within the same half-dimer. Significant differences between network models involved the fragmentation of these five major communities, particularly 2-3a and 3b-4a, denoted by light and dark variations on each color (light and dark blue, green, and orange). A small community in helix 1 was also occasionally observed (yellow).

Dynamical network analysis is commonly applied to study functional dynamics in proteins and can shed light on mechanisms of allosteric communication and molecular signalling [39]. Network models comprise nodes representing residues, and here edges between nodes are based on correlation measured in residue motions within large conformational ensembles. Communities are partitions of the network, within which there are more and stronger connections between nodes. They may be thought of as structural domains, but are defined and grouped purely according to dynamics [36]. Each Cp network model exhibited between seven and eleven communities, with the number of communities identified for a given dimer system varying over the three sub-datasets used for analysis (Figure S9).

Consistently, each model included at least five communities in the body of the dimer, and as many as four in the C-terminal tails. One community typically encompassed helix 1 and 5 for each half-dimer (Figure 4, purple and red). Importantly, helix 5 forms the interdimer interface or contact domain. An additional community typically encompassed helix 2-3a for each half-dimer (Figure 4, blue and green), which comprises a portion of the chassis domain [11]. The remainder of the Cp domain commonly described as the fulcrum [11] was alternately involved in the helix 1,5 or helix 2-3a community of its parent half-dimer, or split between these two communities depending on the sub-dataset. A fifth community typically encompassed helices 3b-4a of both half-dimers in the spike tip (Figure 4, orange), which is also considered a distinct domain [11]. The spike tip is an important aspect of the intradimer interface, and the latter was the only community shown to significantly bridge the two half-dimers. Like the fulcrum, helix 4b could be associated with either the helix 1,5 or 2-3a community. Key communities and their associated Cp domains are summarized in Table 2; Figure 1a provides a reference for the locations and compositions of established Cp domains. Significant differences between network models involved fragmentation of the five major communities.

**Table 2.**
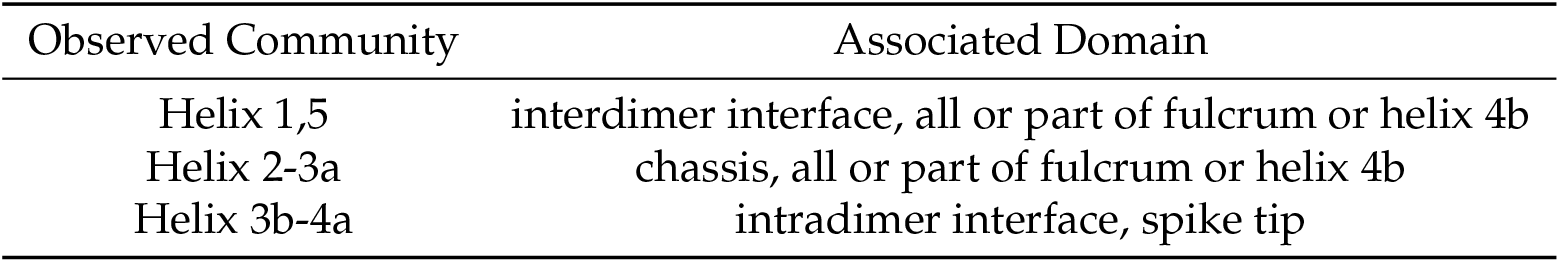
Major communities identified in dynamical network models.

| Observed Community | Associated Domain |
| --- | --- |
| Helix 1,5 | interdimer interface, all or part of fulcrum or helix 4b |
| Helix 2-3a | chassis, all or part of fulcrum or helix 4b |
| Helix 3b-4a | intradimer interface, spike tip |

The helix 1,5 communities are consistently prominent and highly correlated across all network models (Figure 4, purple and red). This may arise in part from interdimer contact and constraints imposed by capsid quaternary structure enforcing shared motions. The concerted displacement of helix 5 during capsid incorporation has been proposed to modulate the assembly process [40]. In one apo-form chain D model, and two AT130-bound chain B models, a small community within helix 1 developed, with minimal attachments to the chassis (Figure 4c and S9, yellow). The loop connecting helix 1 and 2, part of the fulcrum, which has been suggested to mechanically coordinate the motions of the interdimer interface and the spike [11], showed the ability to participate in the helix 1,5 community through weakly correlated motions (Figure 4).

Distinct C-terminal tail communities found in the apo-form system were absorbed by helix 1,5 communities in 4 out of 6 models of the AT130-bound system (Figure S9). That contact of the CpAM can establish communication between helix 5 and the C-terminals of Cp149 (which serve as flexible linkers to the RNA-binding CTDs in full-length Cp183 [41]) raises the possibility of a secondary mechanism underlying AT130’s ability to disrupt the process of genome packaging, apart from mistiming. On the other hand, all network models of AT130-bound chain D show the C-terminal tail community encroaching into that of helix 5 (Figure 4d, red), as far up the sequence as residue 126, likely resulting from direct interaction with the CpAM. Importantly, ordering of residues 126-136 has been correlated with capsid formation [40,42]. Secondary structure analysis showed that a 310 helix in this region becomes less likely in the A and D chains, yet more likely in the B chain in the presence of AT130 (Figure 3).

The helix 2-3a communities and the remainder of the chassis represent the regions of Cp with the least degree of correlated motion (Figure 4). Nevertheless, their dynamics are affected by AT130 in unexpected ways. Although it manifests as a single community in all models of apo-form chain A, the helix 2-3a community splits into separate helix 2 and helix 3a communities in all AT130-bound models (Figure 4a,c and S9, light and dark green). This is an allosteric effect, possibly arising from changes in capsid morphology [10], as chain A does not contact the CpAM. Separate helix 2 and helix 3a communities are observed in two out of three models of chain B, regardless of direct interaction with AT130 (Figure 4a,c and S9, light and dark blue). Helix 2 and helix 3a of chain C are split into separate communities in two out of three models in the apo-form system, but only one out of three models in the AT130-bound system (Figure 4b,d and S9, light and dark green), which likewise involves direct contact. A single helix 2-3a community is found in all models of chain D regardless of the CpAM (Figure 4b,d and S9, light and dark blue). Interestingly, chains A and D, which consistently exhibit combined helix 2-3a communities in the absence of AT130, also represent the quasi-equivalent pockets that are not amenable to AT130 binding [10].

The helix 3b-4a community is consistently prominent across all network models (Figure 4, orange). The strength of dynamical correlation within the community is strong when the probability of helicity is high, and is clearly decreased for the case of the apo-form CD dimer where the majority of spike tips remain only partially-folded (Figure 4b). Reduced helicity in the spike tips has been shown to disrupt the intradimer interface and negatively impact assembly [16]. Remarkably, in one model of the AT130-bound CD dimer, the helix 3b-4a community splits into two separate communities, one for each half-dimer (Figure 4b and S9, light and dark orange), indicating diminished communication across the interface. Correlation within the individual half-dimer spike tips is also weakened. The structural origin of this tertiary disruption in the presence of AT130 can be attributed to dramatic bending of the dimer hinge (discussed in the following section), which results in an open spike conformation and broken intradimer interface with an increased probability in chain D.

### 3.3. Hinge Curvature Analysis

Figure 5 shows distributions of curvature adopted by the dimer hinge for the AB and CD capsid-incorporated dimers in the apo-form and AT130-bound systems during MD simulations. Importantly, Cp contains a hinge that connects the spike tip and chassis domain. The hinge is defined by conserved Gly63 and Gly94, which impart a bending compliance to helix 3 and 4, respectively, and partition them into helices 3a/b and 4a/b [11]. We have previously used MD simulations to characterize hinge bending in free dimers, showing that fully-folded spike tips exhibited smaller angles of helix curvature and greater potential for a closed intradimer interface [16]. Here, the analysis is repeated for capsid-incorporated dimers, measuring the angle at the point of highest curvature along helix 4 (the hinge vertex), which was consistently found to be Val93 in both free and capsid-incorporated dimers. Note that the reported angles capture helix 4’s deviation from linearity resulting from bending of the hinge; larger angles indicate more bending.

**Figure 5.**
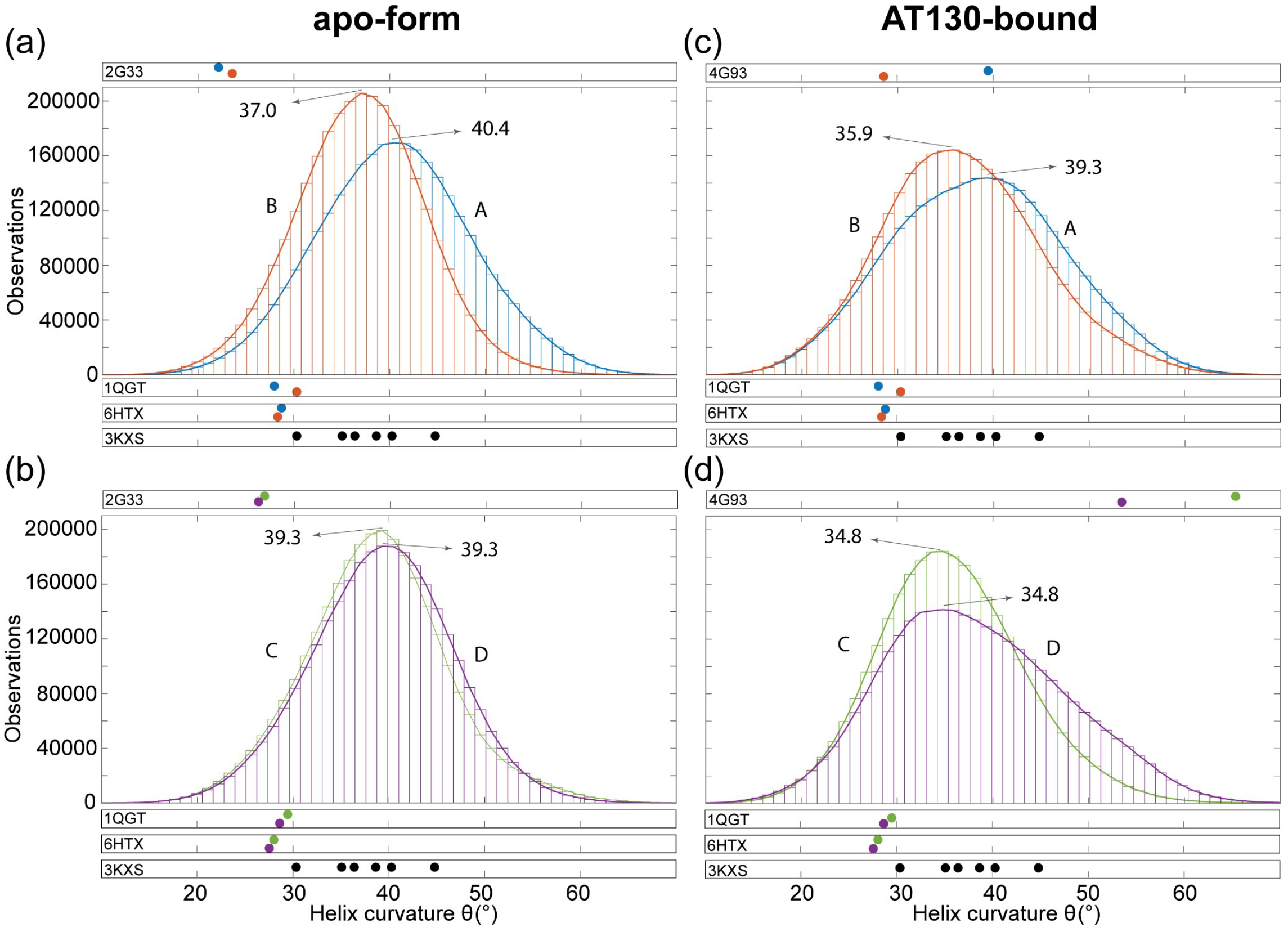
Hinge curvature analysis of capsid-incorporated dimers. (**a-b**) Distribution of angles for apo-form AB and CD dimers. (**c-d**) Distribution of angles for AT130-bound AB and CD dimers. Quasi-equivalent chains A, B, C, and D are colored blue, orange, green, and purple, respectively. Angles denote the curvature or deviation from linearity at the hinge vertex (Val93), measured with Bendix [37] along residues 88-98. Distributions calculated based on ensembles of three million conformers collected over 60 *µ*s of cumulative sampling. Histograms use a bin size of 80. Centers of distributions are indicated, and angles measured for relevant experimental structures are provided for reference.

All four quasi-equivalent chains exhibit significantly higher average hinge angles than those observed in experimental structures of the apo-form capsid [4,32,38], more similar to those observed in crystals of assembly-incompetent dimers [11] (Figure 5). This result is noteworthy, as comparison of capsid and dimer structures has inspired hypotheses that relaxed hinges and closed intradimer interfaces are a feature of assembly-active Cp conformations [16]. MD simulations indicate that capsid-incorporated and free dimers are more alike in spike conformation than experiments suggest. On the other hand, hinge angles measured for the AT130-bound crystal model [10], particularly in the CD dimer, are quite extreme (Figure 5d). Importantly, these values are artifacts arising from highly disordered secondary structure within the hinge region, which leads to inaccurate fitting of helix abstractions and assessment of curvature [37].

The distributions of curvature measured for MD simulations of the apo-form capsid are Gaussian for all four quasi-equivalent chains (Figure 5a-b). The A chain distribution is markedly flatter than the other three. As the A chain forms capsid pentamers, quasi-equivalence likely accounts for differences in hinge behavior relative to the B, C, and D chains that form capsid hexamers. The A chain distribution, centered at 40.4^*°*^, also exhibits the highest angle (most dramatically bent hinge), while the B chain distribution, centered at 37.0^*°*^, exhibits the lowest (Figure 5a). Both A and B spike tips are well-folded during MD simulations, yet well-folded free dimers exhibited distributions centered at 37.0*°*(based on data from [16]), indicating that preferred hinge states are influenced by quasi-equivalence and constraints imposed by capsid quaternary structure. It is likely that because the A chain hinge is more bent on average, the B chain must necessarily be less bent on average to maintain a closed intradimer interface. An example of an apo-form AB dimer sampled during MD simulations is shown in Figure 6a.

**Figure 6.**
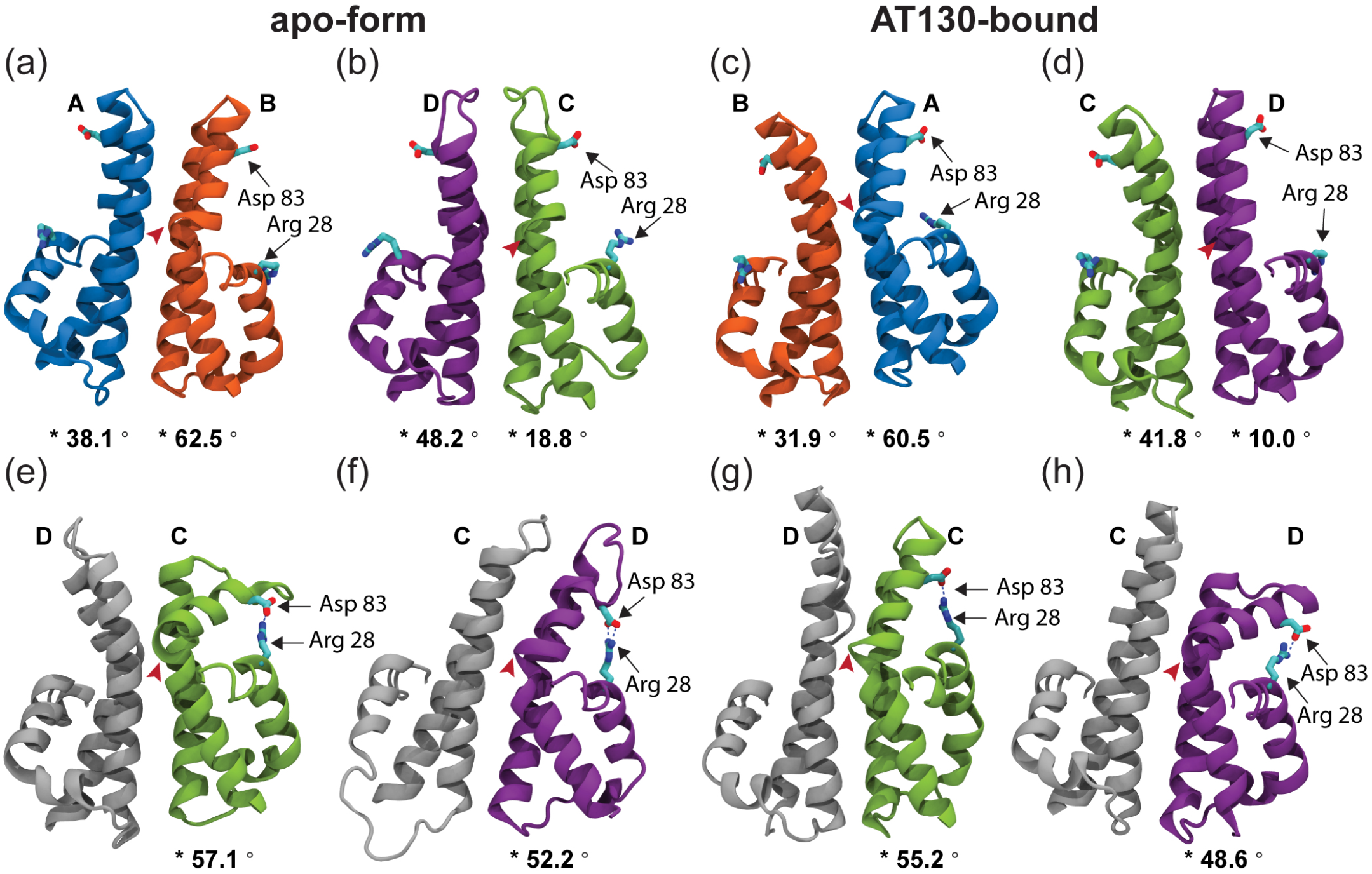
Notable conformations of capsid-incorporated dimers. (**a-b**) Representative structures for apo-form AB and CD dimers. (**c-d**) Representative structures for AT130-bound AB and CD dimers. Quasi-equivalent chains A, B, C, and D are colored blue, orange, green, and purple, respectively. Arg28 and Asp83, which can form a rare salt bridge, are labeled. Apex of the dimer hinge (Val93) is denoted by a red arrow and measurement of curvature at the apex is given beneath. (**e-h**) Structures for apo-form and AT130-bound CD dimers illustrating conformations that allow the Arg28-Asp83 salt bridge, enabling direct contact between the spike and fulcrum. All structures are aligned based on the dimer chassis. Conformations with large hinge angle and intact salt bridge within a half-dimer exhibit an open spike tip and disrupted intradimer interface. This state was observed most often for the AT130-bound D chain (Figure 10).

Interestingly, the C and D chain distributions are balanced in average hinge angle, both centered at 39.3*°*(Figure 5b). The C and D spike tips have a high probability to be only partially-folded during MD simulations, and partially-folded free dimers exhibited distributions also centered at 39.3*°*(based on data from [16]). This result indicates that secondary structure is the primary determining factor for hinge behavior in CD dimers in the absence of AT130. The Glu77-Arg82 salt bridge, which was observed frequently in MD simulations of the apo-form CD dimer, can cause larger average hinge angles by stabilizing a loop at the apex of the partially-folded spike tip that has the potential to protrude into the interface, leading to steric clash and electrostatic repulsion that is alleviated by adopting increased curvature along helix 4 [16]. An example of an apo-form CD dimer sampled during MD simulations is shown in Figure 6b.

The distributions of curvature measured for MD simulations of the AT130-bound capsid are shifted and flattened for all four quasi-equivalent chains compared to apo-form (Figure 5c-d). Further, the distributions are only strictly Gaussian for B and C chains, which contain the occupied CpAM binding pockets, while the distributions for A and D chains exhibit distortion. Higher populations of above- and below-average hinge angles in the AT130-bound capsid could partially account for lower resolution in the crystal model [10], a consequence of averaging over more pronounced conformational variation in spikes. Importantly, the spike tips of all four quasi-equivalent chains are well-folded during MD simulations, so changes in hinge behavior are not a consequence of disordered secondary structure, but arise from the presence of AT130. The A and B chain distributions are shifted down by exactly 1.1*°*(Figure 5c), while the C and D chain distributions are shifted down by exactly 4.5*°*(Figure 5d), indicating relaxation of the hinge on average. An example of an AT130-bound AB and CD dimer sampled during MD simulations is shown in Figure 6c-d. Notably, the D chain of AT130-bound dimers exhibits an unusual propensity for pronounced hinge bending during MD simulations, accounting for distortion on the right-side of the distribution (Figure 5d). Upon examination, it was discovered that larger hinge angles in the D chain often lead to marked opening of the spike tip and breaking of contact across the intradimer interface. This behavior explains fragmentation of network communities in the spike tip in the presence of AT130. Further, this open conformation facilitates formation of a salt bridge between Arg28-Asp83, which introduces a direct contact between the spike tip and fulcrum and provides a mechanistic explanation for allosteric communication across these domains (Figure 6e-h).

Rarely, the open conformation, along with the salt-bridge, was observed in apo-form CD dimers (Figure 6e), but never in AB dimers with or without AT130. Interestingly, the presence of the salt bridge is not necessarily an indication of an open spike conformation, as disorder in helix 4a or displacement of helix 2 can bring Asp28 in proximity to interact with Asp83 without breaking the intradimer interface (Figure 6f-g). Disruption in the chassis was possible, by prying away helix 4b, but network analysis did not indicate a role for the base of the four-helix bundle in intradimer communication. A partially-folded helix 3b appears to increase opportunity for the salt bridge. Overall, the Arg28-Asp83 interaction occurred with low probability, but was observed more often in the D chain with AT130 (Figure 6h). Time series of hinge angles and Arg28-Asp83 contact for apo-form and AT130-bound CD dimers are provided in Figure S10.

### 3.4. Principal Component Analysis

Figure 7 shows the top five modes of essential dynamics observed for the AB and CD capsid-incorporated dimers in the apo-form and AT130-bound systems based on principal component analysis of MD simulations. Our previous work on the apo-form capsid examined essential dynamics for the complete icosahedron, reporting asymmetric distortion and complex collective motions of dimers within the capsid surface [12]. Overall, the intact capsid modes proved difficult to interpret due to limited conformational sampling.

**Figure 7.**
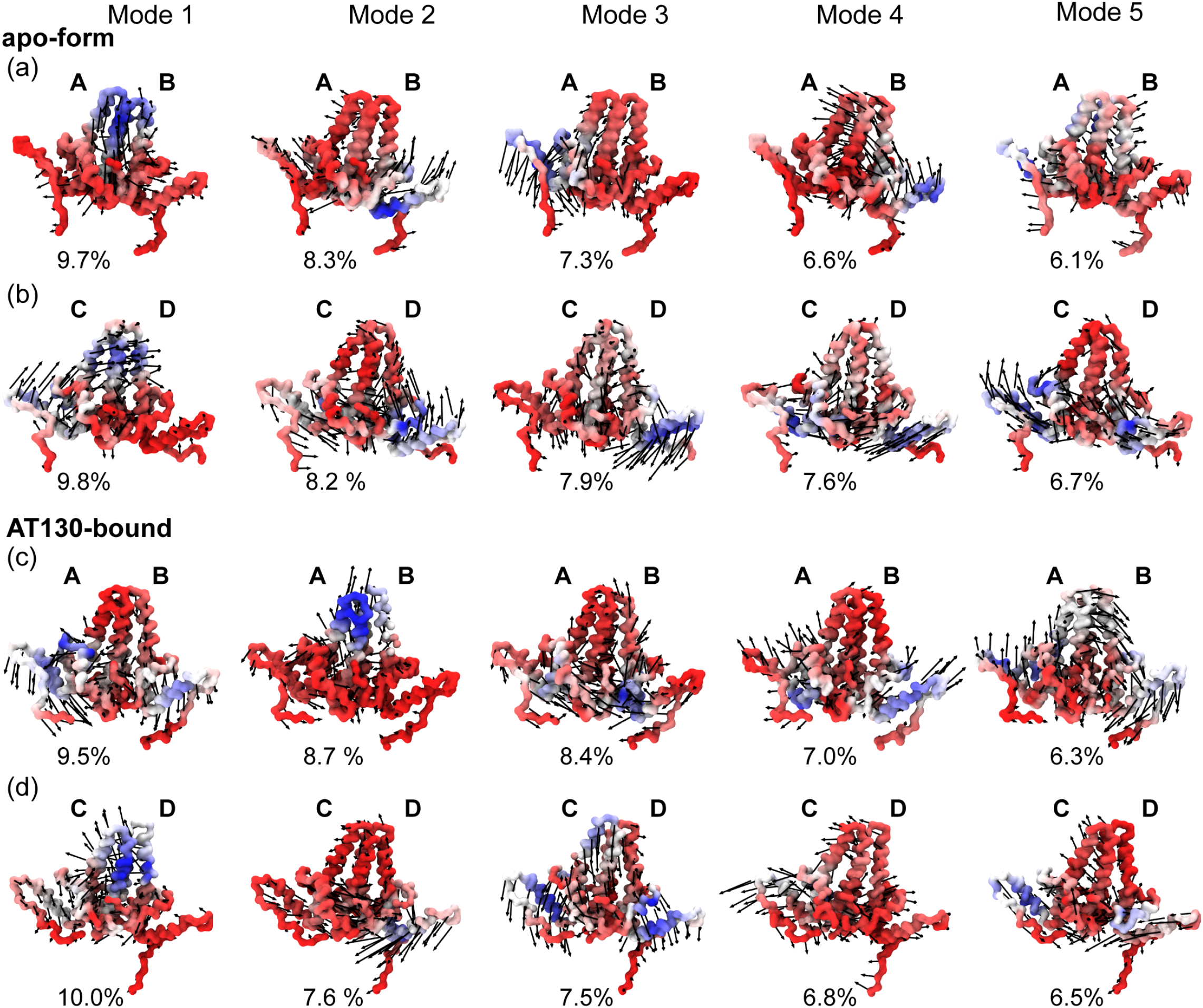
Essential dynamics of capsid-incorporated dimers. (**a**) Apo-form dimer AB. (**b**) Apo-form dimer CD. (**c**) AT130-bound dimer AB. (**d**) AT130-bound dimer CD. Backbone trace of dimers shown for each of the top five modes of essential dynamics, calculated based on principal component analysis of 60 *µ*s of cumulative simulation sampling. Blue to red color scale denotes high to low mobility. Black arrows indicate direction of motion and are scaled by 12 for visual clarity. Percentages indicate contributions to variance, which total 38.0%, 40.2%, 39.9%, and 38.4%, respectively for panels **(a-d)**.

Here, each capsid-incorporated dimer system benefits from 60× more sampling, providing the advantage of well-converged datasets.

Twenty modes were calculated per dimer, and for each system, the top five modes together account for ~40% of total variance. The first (lowest frequency) modes consistently contribute 10% or less to variance, ultimately indicating that there is no dominant fluctuation to be distilled from the dynamics. Cp remains a marvel of flexibility, even when constrained by the quaternary structure of the capsid. Due to the diversity of observed motions, it is unclear, even given such a large conformational ensemble, whether the differences in modes across systems arise strictly from quasi-equivalence and the influence of AT130, or if the system remains yet undersampled.

All calculated modes necessarily exhibit motion relative to the chassis, which has been described as the underlying constant substructure around which other dimer domains hinge [11], and was used here as the standard for alignment. Indeed, the majority of mode movements can be linked to distinct domains, including the fulcrum (helix 1-2), spike tip (helix 3b-4a), and interdimer contact domain (helix 5) [11]. Overall, most motions capture flexing of the spike and radial movements of the fulcrum and helix 5; the latter may be related to the ability of the fivefold vertices of the capsid to protrude and sixfold/threefold vertices to flatten, a known morphological adjustment that can accommodate the presence of CpAMs [10,15,32,43].

For the apo-form AB system (Figure 7a), Mode 1 is characterized by spike waving, primarily on the side of chain A, consistent with the observation that its average dimer hinge angle is larger than that of chain B; helix 3 of chain A concomitantly sinks into the dimer core. Mode 2 is characterized by a slight twisting of the half-dimers against each other at the chassis, along with upward motion of the fulcrum and helix 5 of chain B. Mode 3 is the converse of this movement, characterized by downward motion of the fulcrum and helix 5 of chain A, with corresponding motion of the spike. Mode 4 shares similarities with Mode 2, but the movement of the fulcrum of chain B is skewed toward chain A and carries the spike with it. Mode 5 involves a lateral stretching of helix 5 of chain A and opposite waving of the spike.

For the apo-form CD system (Figure 7b), Mode 1 is characterized by concerted motion of chain C, along with the spike tip of chain D, consistent with the observation from network models that helix 3b-4a is the key location of intradimer communication; helix 5 of chain C undergoes an upward motion, perhaps shifting along with chain B where the two share an interface. Mode 2 is characterized by a rotary motion at the chassis, which sends the fulcrum and helix 5 of chain D upward. Mode 3 shares similarities with Mode 2 of the AB dimer, with slight twisting of the half-dimers against each other that sends the fulcrum and helix 5 of chain B downward. Mode 4 involves a lateral compression of the dimer, particularly along helix 4b-5 of chain D. Mode 5 pushes the fulcrum and helix 5 of chain C upward.

For the AT130-bound AB system (Figure 7c), Mode 1 is characterized by a slight twisting of the half-dimers against each other at the chassis, and a downward motion of the fulcrum and helix 5 of both chains. Mode 2 is characterized by spike movement, particularly in chain A; interestingly the spike tips do not travel in the same direction. Mode 3 involves rotation of the half-dimers toward each other at the chassis, pressing upward the spike tip of chain B. Mode 4 shares similarities with Mode 1, entailing a dimer expansion that pushes the fulcrum and helix 5 of both chains upward, and that of chain B to the side. Mode 5 is a rotation of the dimer in the direction of chain B, which causes a seesawing movement along the helix 5 axes.

For the AT130-bound CD system (Figure 7d), Mode 1 is characterized by spike movement, particularly in chain C, similar to Mode 2 of the AB dimer; again the spike tips do not travel in the same direction. Mode 2 is characterized by a familiar downward motion of helix 5 of chain D. Mode 3 involves concerted downward motion of the entire dimer, except the spike tip of chain D, which may arise from the conformations that exhibit a broken intradimer interface with disrupted cross-communication. Mode 3 entails a pressing of helix 5 of chain C downward. Mode 5 includes a contraction of helix 5 of both chains inward toward the chassis.

## 4. Discussion

Here, all-atom MD simulations were applied to study the HBV capsid in complex with the phenylpropenamide AT130 [10]. Comparison with the apo-form HBV capsid [32] based on an analogous, previously obtained MD simulation [12], enabled detection of changes in structure and dynamics induced by the bound CpAM. Importantly, the investigation of intact capsids on microsecond timescales afforded 60 *µ*s of cumulative sampling for each quasi-equivalent capsid-incorporated dimer, providing an unprecedented conformational ensemble for the characterization of Cp behavior within the context of native physiological conditions. These data have proved essential for revealing the true nature of the AT130-bound HBV capsid, beyond what has been possible with experimental structures and other biophysical studies.

While a crystal model of the capsid complexed with AT130 reported significant disruption of secondary structure, particularly in helix 3b-4a of the spikes [10], MD simulations based on this model showed that the majority of spikes rapidly refolded (Figure 3). Spikes exhibited a reduced ability to refold in the absence of AT130, indicating that, rather than damaging secondary structure, the CpAM facilitates recovery of spike helicity allosterically. Previous work has shown that mutations in the spike that impair refolding of helix 4a have a negative impact on capsid assembly [16]. It follows that maintenance of secondary structure in helix 4a is an aspect of AT130’s assembly accelerating mechanism. Well-folded spike helices are associated with a stable intradimer interface, which has been linked to efficient assembly [44].

While a large body of evidence supports the existence of an allosteric connection between the intra- and interdimer interfaces, and that CpAMs tap into this allostery to speed oligomerization, the atomistic details of Cp’s allosteric network have remained elusive. Experiments indicate that the network spans the entire protein, entails concerted dynamics, and involves systematic movements of distinct dimer domains [11,42,44,45]. Network models based on MD simulations confirm that previously established dimer domains share strongly correlated dynamics and that adjustments occur with respect to a central chassis (Figure 4). When spike tips are well-folded, dynamical cross-talk between half-dimers is prominent. Helix 5, the interdimer interface that mediates subunit contact, can communicate directly with the fulcrum through correlated motion. Helix 4b of the chassis can participate in their same network community, and may, thus, play a role in transducing signal between the intra- and interdimer interfaces via its covalent connection to helix 4a and tertiary relationship with the fulcrum.

Alternatively, a new conformational state revealed here by MD simulations allows for direct contact between the spike tip and fulcrum via an Arg28-Asp83 salt bridge. Observation of this relatively rare state was made possible only by long-timescale examination of many copies of Cp. Formation of the salt-bridge often occurred at the expense of the intradimer interface, requiring either unfolding of the spike tip or severing of association of half-dimers along helix 3b-4a; however, some conformers with the intact salt-bridge retained intradimer communication, particularly if adjustments were made in positioning of the fulcrum or helix 4b (Figure 6f-g).

Such adjustments are likely more feasible in free dimers than in capsid-incorporated ones constrained by quaternary structure, such that the Arg28-Asp83 salt bridge may represent a literal missing link in the Cp allosteric mechanism. When present, the salt bridge facilities direct communication between the spike tip (helix 4a) and fulcrum (helix 2), which is linked dynamically with the contact domain (helix 5), thus, connecting the intra- and interdimer interfaces. Even fleeting contact may be sufficient to transduce signal. Importantly, both Arg28 and Asp83 are highly conserved residues, appearing in 99% and 100% of deposited Cp sequences [46]. Indeed, the 1% mismatch refers to an R28K substitution, where lysine would be similarly capable of forming the salt bridge.

With respect to the role of CpAMs in modulating this allostery, AT130 physically contacts helix 2 and 5, which network models show can communicate via shared motions within the fulcrum loop. The CpAM also showed the ability to induce helical secondary structure in residues 126-136, depending on quasi-equivalence, as well as dynamical correlation between this region and the C-terminal tails. Ordering of residues 126-136 has been associated with capsid assembly [40,42], and a key aspect of AT130’s antiviral mechanism is the acceleration of assembly such that genome, which is bound by the C-terminal linked CTDs, is not encapsulated [7,8]. The latter suggests that AT130 may be able to communicate with the CTDs allosterically via these correlated motions that extend from its binding site down the Cp149 tails, which may influence RNA binding and contribute to the mistiming of genome packaging.

Comparison of capsid and dimer crystals has inspired hypotheses that closed intradimer interfaces are a feature of assembly-active Cp conformations [16]. Structures previously described as open [11,16] pale in comparison to rare conformations observed during MD simulations that exhibit notable bending of the dimer hinge and broken contact between half-dimers in the spike tip. As evidenced by network models, even a relatively small population of these structures was sufficient to diminish communication between half-dimers and affect the allostery spanning Cp. Such open and closed states are likely still related to the conformational switch that activates capsid assembly [11,42,45], but the discovery of the Arg28-Asp83 salt bridge raises new possibilities for the mechanism of switching. Mutations in the intradimer interface that alter assembly kinetics support an allosterically regulated switch. It may be that crosslinking of dimers at Cys61, which slows assembly [47], inhibits formation of the salt bridge, or that the F97L substitution, which speeds assembly [48], facilitates adjustment of helix 4b to accommodate the salt bridge without compromising intradimer contact.

The open dimer conformation that allows the Arg28-Asp83 salt bridge, as well as higher overall populations of above- and below-average dimer hinge angles (Figure 5) could partially account for low resolution obtained for spike tips in the AT130-bound crystal model, particularly in the D chain, whose secondary structure was reported as disordered all the way down to the hinge vertex [10]. The hinge contributes significantly to spike flexibility. Notably, principal component analysis emphasises that some spike motion is also possible orthogonal to the direction of hinging (Figure 7). It may be that binding of CpAMs in B and C sites (which are capped, respectively, by C and D chains) somehow reduces quaternary stress on the capsid to facilitate spike refolding and allow for increased spike flexibility. As dimer hinge angles observed in crystal structures were not representative of the distributions captured in MD simulations, these structures likely do not adequately represent the capsid’s native state.

Our previous work has shown that variability in the conformational ensemble accounts for poor resolution in most experimentally-derived HBV capsid structures, a consequence of averaging over diverse ensembles describing a highly flexible protein [12]. In this case, it seems that low crystallographic resolution arising from variation in spike conformations was interpreted as distorted secondary structure, leading to erroneous conclusions about the structural effects of AT130. This outcome underscores the limitations of experimental methods and the insufficiency of a single conformation for explaining structure-function relationships in complex biomolecular systems. Importantly, MD simulations can reveal details that elude other structural biology techniques, and represent an essential complement to experiments in the study of virus capsids and their interactions with small molecule inhibitors and cellular host factors [13,15].

## Author Contributions

J.A.H.-P. designed the project, constructed models, performed simulations, secured funding and resources, and trained students. B.C.G. carried out AT130 parameterization. C.P.S. performed data analysis and produced figures. All authors contributed to the preparation of the manuscript. All authors have read and agreed to the published version of the manuscript.

## Funding

Research reported in this publication was supported by the National Institutes of Health under award P20-GM-104316 and the National Science Foundation under award MCB-2027096, funded in part by Delaware Established Program to Stimulate Competitive Research. This research is part of the Blue Waters sustained-petascale computing project, which is supported by the National Science Foundation (awards OCI-0725070 and ACI-1238993) and the state of Illinois. Blue Waters is a joint effort of the University of Illinois at Urbana-Champaign and its National Center for Supercomputing Applications. This research is part of the Frontera computing project at the Texas Advanced Computing Center. Frontera is made possible by the National Science Foundation under award OAC-1818253.

## Data Availability Statement

The dataset that supports the findings of this manuscript is available from the corresponding author upon request.

## Conflicts of Interest

The authors declare no conflict of interest. The funders had no role in the design of the study; in the collection, analyses, or interpretation of data; in the writing of the manuscript, or in the decision to publish the results. The content is solely the responsibility of the authors and does not necessarily represent the official views of the National Institutes of Health.

## Abbreviations

The following abbreviations are used in this manuscript:

Cp: Core protein
Cp149: Core protein assembly domain, residues 1 to 149
Cp183: Full-length core protein, residues 1 to 183
CpAM: Core protein allosteric modulator
CTD: C-terminal domain
HBV: Hepatitis B virus
MD: Molecular dynamics
PPA: Phenylpropenamide
PDB: Protein Data Bank QM Quantum mechanics

## 5. Supplementary Figures

**Figure 8.**
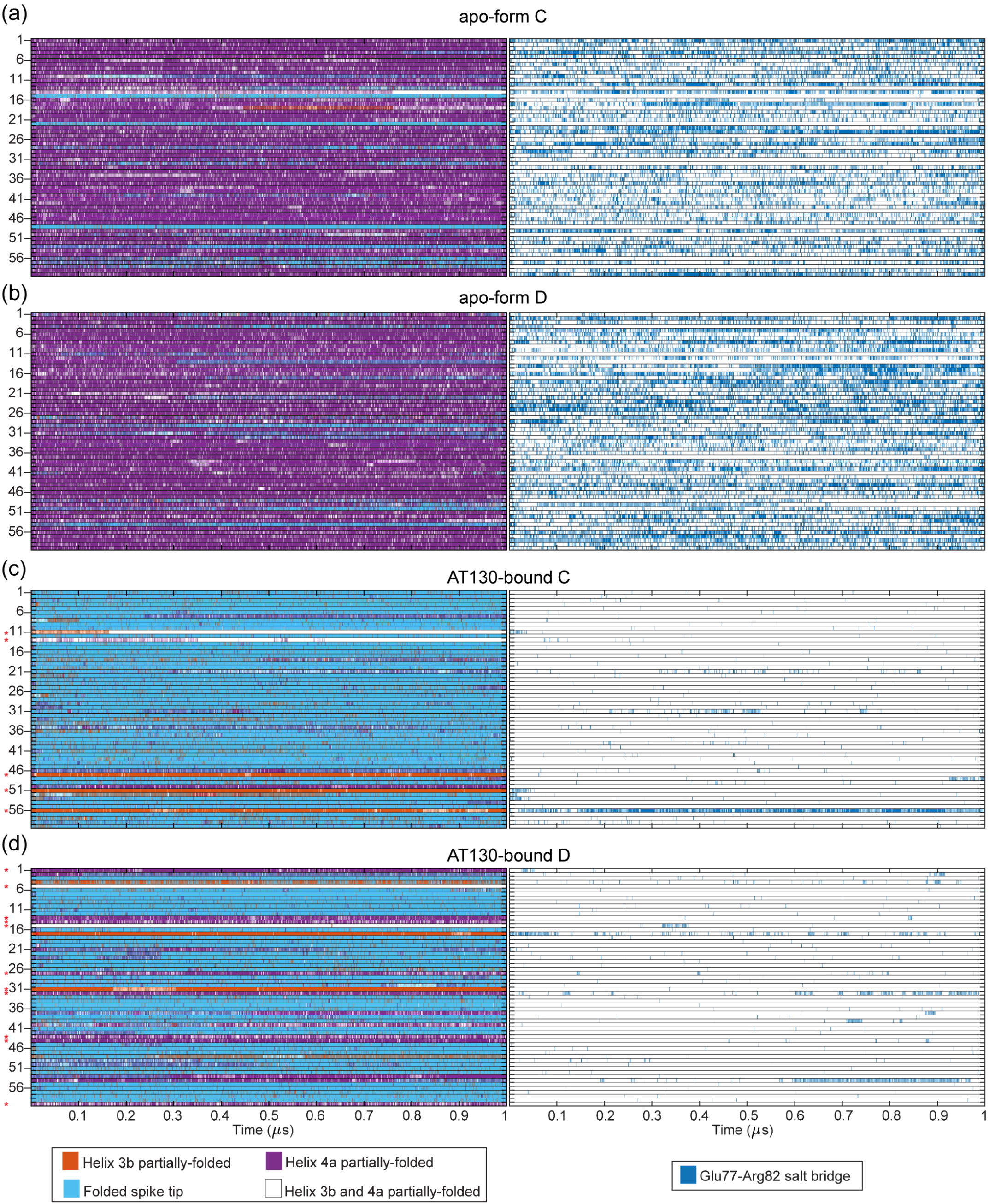
Spike helicity and Glu77-Arg82 salt bridge. Heat maps showing time series of spike tip helicity (left) and presence of Glu77-Arg82 salt bridge (right) for apo-form (**a-b**) and AT130-bound (**c-d**) CD dimers. Fully-folded spike tip is defined as having helix 3b intact through its C-terminus at Asn75 and helix 4a intact from its N-terminus at Pro79; less helical content is considered partially-folded. The Glu77-Arg82 salt bridge has been shown to stabilize partially-folded spike conformations [16], and its common occurrence here likely relates to the reduced ability of apo-form spikes to recover helicity.

**Figure 9.**
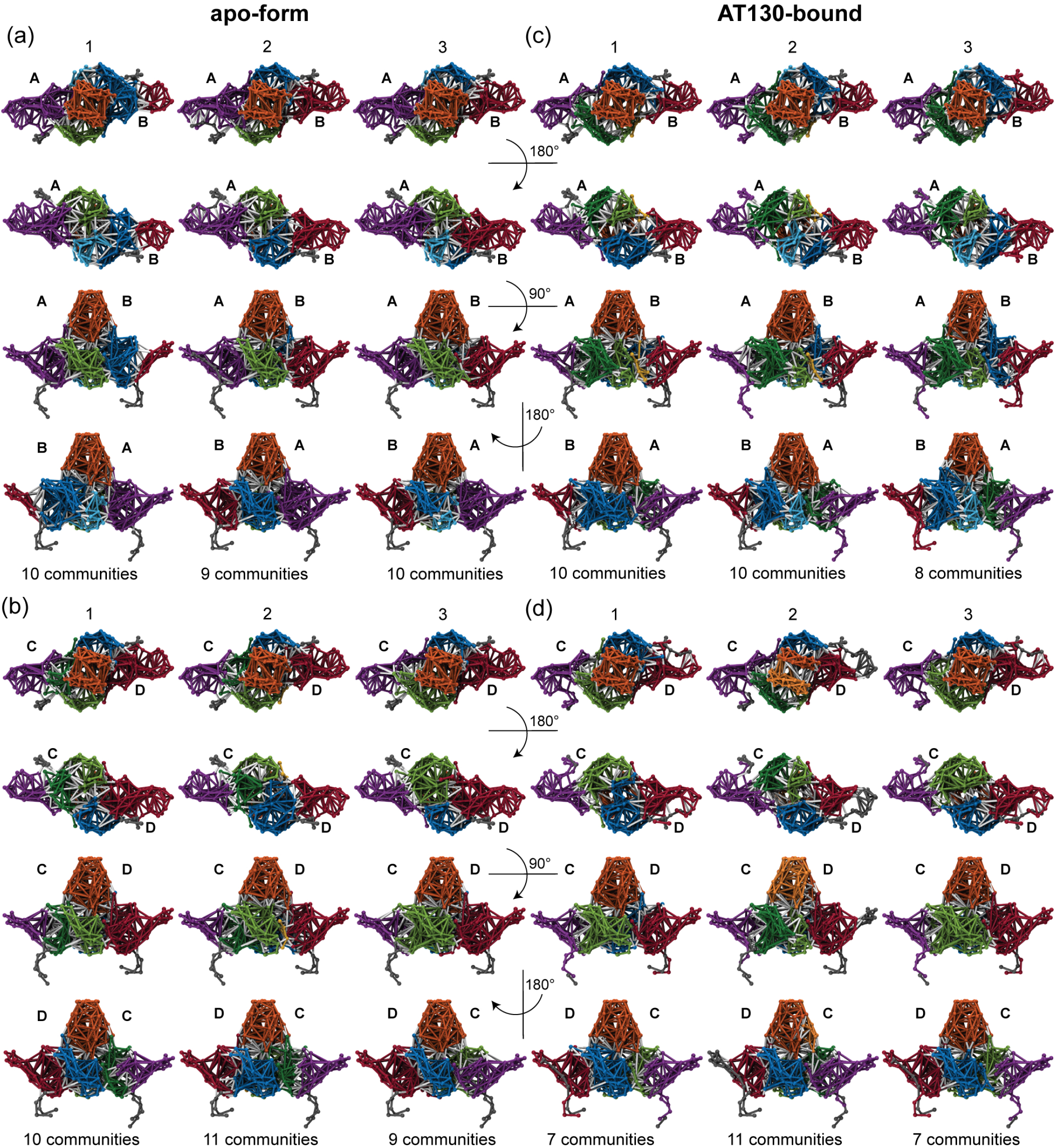
Complete set of twelve network models. Dynamical network models calculated in triplicate for apo-form (**a-b**) and AT130-bound (**c-d**) dimers. Each model based on an ensemble of one million randomly-selected conformers collected over 20 *µ*s of cumulative sampling. Colors denote sub-network communities. Five major communities encompass helix 1,5 of each half-dimer (purple and red), where helix 5 forms the interdimer interface, helix 2-3a of each half-dimer (blue and green), which are part of the chassis domain, and helix 3b-4a of both half dimers (orange), called the spike tip. The remainder of the fulcrum domain participated in either the helix 1,5 or helix 2-3a community within the same half-dimer. Significant differences between network models involved the fragmentation of these five major communities, particularly 2-3a and 3b-4a, denoted by light and dark variations on each color (light and dark blue, green, and orange). A small community in helix 1 was also occasionally observed (yellow).

**Figure 10.**
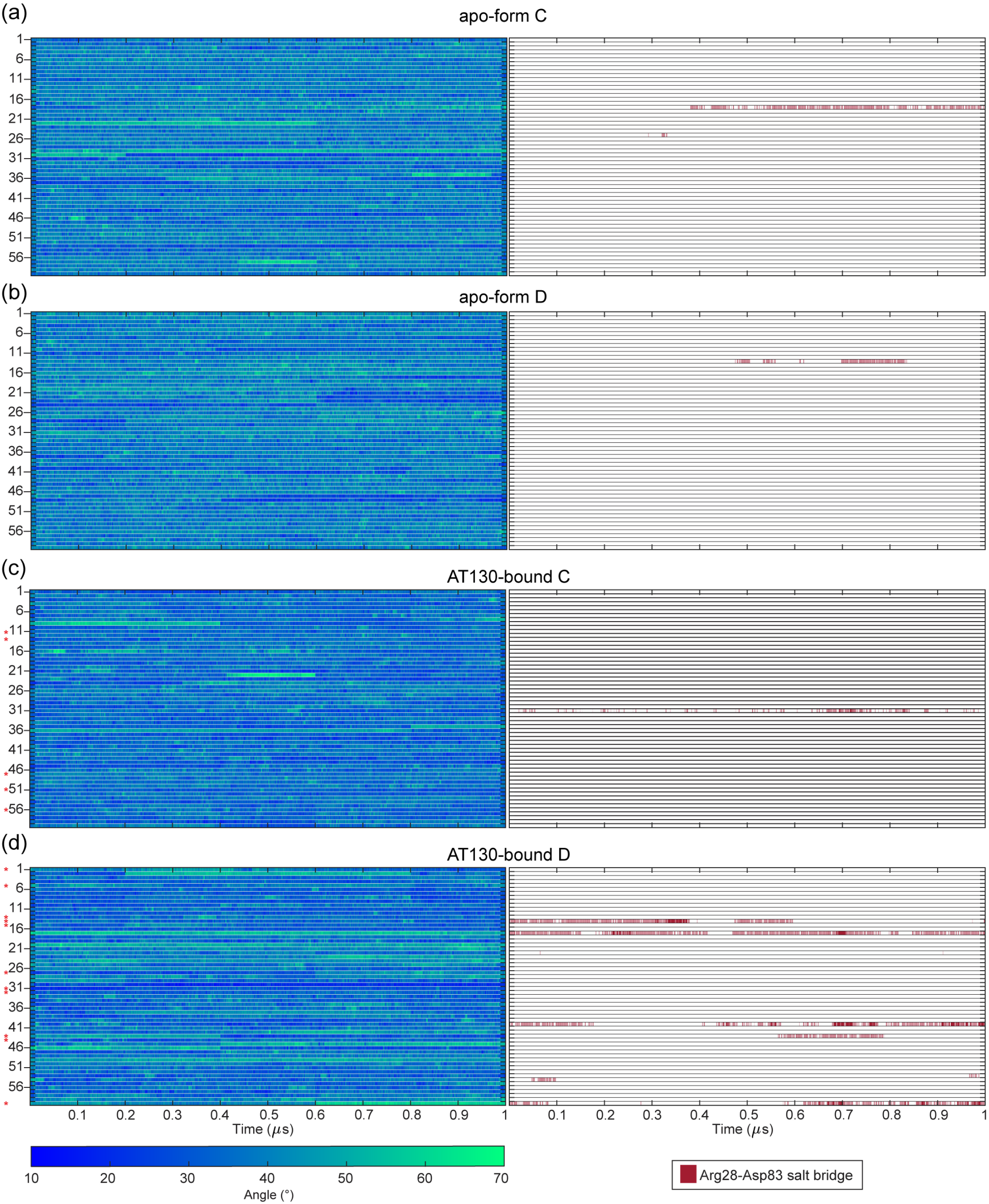
Hinge angle and Arg28-Asp83 salt-bridge. Heat maps showing time series of hinge angle (left) and presence of Arg28-Asp83 salt-bridge (right) for apo-form (**a-b**) and AT130-bound (**c-d**) CD dimers. Salt bridge interaction is typically correlated with dramatic curvature along helix 4, facilitating residue contact between the spike tip and fulcrum.

## Notes

### Competing Interest Statement

The authors have declared no competing interest.

## References

1. Venkatakrishnan, B., Zlotnick, A. The structural biology of hepatitis B virus: Form and function. Annual Review of Virology 2016, 3, 429–451.

2. Razavi-Shearer, D., Gamkrelidze, I., Nguyen, M.H., Chen, D.S., Van Damme, P., Abbas, Z., Abdulla, M., Abou Rached, A., Adda, D., Aho, I., others. Global prevalence, treatment, and prevention of hepatitis B virus infection in 2016: A modelling study. The Lancet Gastroenterology &Hepatology 2018, 3, 383–403.

3. Organization, W.H. Global health sector strategy on viral hepatitis 2016-2021. Towards ending viral hepatitis. Technical report, World Health Organization, 2016.

4. Wynne, S., Crowther, R., Leslie, A. The crystal structure of the human hepatitis B virus capsid. Molecular Cell 1999, 3, 771–780.

5. Schlicksup, C.J., Zlotnick, A. Viral structural proteins as targets for antivirals. Current Opinion in Virology 2020, 45, 43–50.

6. Perni, R.B., Conway, S.C., Ladner, S.K., Zaifert, K., Otto, M.J., King, R.W. Phenylpropenamide derivatives as inhibitors of hepatitis B virus replication. Bioorganic &Medicinal Chemistry Letters 2000, 10, 2687–2690.

7. Feld, J., Colledge, D., Sozzi, V., Edwards, R., Littlejohn, M., Locarnini, S. The phenylpropenamide derivative AT-130 blocks HBV replication at the level of viral RNA packaging. Antiviral research 2007, 76, 168–177.

8. Katen, S.P., Chirapu, S.R., Finn, M., Zlotnick, A. Trapping of hepatitis B virus capsid assembly intermediates by phenylpropenamide assembly accelerators. ACS Chemical Biology 2010, 5, 1125– 1136.

9. Kondylis, P., Schlicksup, C.J., Katen, S.P., Lee, L.S., Zlotnick, A., Jacobson, S.C. Evolution of intermediates during capsid assembly of hepatitis B virus with phenylpropenamide-based antivirals. ACS Infectious Diseases 2019, 5, 769–777.

10. Katen, S.P., Tan, Z., Chirapu, S.R., Finn, M., Zlotnick, A. Assembly-directed antivirals differentially bind quasiequivalent pockets to modify hepatitis B virus capsid tertiary and quaternary structure. Structure 2013, 21, 1406–1416.

11. Packianathan, C., Katen, S.P., Dann, C.E., Zlotnick, A. Conformational changes in the hepatitis B virus core protein are consistent with a role for allostery in virus assembly. Journal of Virology 2010, 84, 1607–1615.

12. Hadden, J.A., Perilla, J.R., Schlicksup, C.J., Venkatakrishnan, B., Zlotnick, A., Schulten, K. All-atom molecular dynamics of the HBV capsid reveals insights into biological function and cryo-EM resolution limits. eLife 2018, 7, e32478.

13. Hadden, J.A., Perilla, J.R. All-atom virus simulations. Current Opinion in Virology 2018, 31, 82–91.

14. Ruan, L., Hadden, J.A., Zlotnick, A. Assembly properties of hepatitis B virus core protein mutants correlate with their resistance to assembly-directed antivirals. Journal of Virology 2018, 92.

15. Perilla, J.R., Hadden, J.A., Goh, B.C., Mayne, C.G., Schulten, K. All-atom molecular dynamics of virus capsids as drug targets. The Journal of Physical Chemistry Letters 2016, 7, 1836–1844.

16. Zhao, Z., Wang, J.C.Y., Segura, C.P., Hadden-Perilla, J.A., Zlotnick, A. The integrity of the intradimer interface of the Hepatitis B Virus capsid protein dimer regulates capsid self-assembly. ACS Chemical Biology 2020.

17. Watanabe, G., Sato, S., Iwadate, M., Umeyama, H., Hayakawa, M., Murakami, Y., Yoneda, S. Molecular dynamics simulations to determine the structure and dynamics of hepatitis B virus capsid bound to a novel anti-viral drug. Chemical and Pharmaceutical Bulletin 2016, 64, 1393–1396.

18. Tu, J., Li, J.J., Shan, Z.J., Zhai, H.L. Exploring the binding mechanism of Heteroaryldihydropyrimidines and Hepatitis B Virus capsid combined 3D-QSAR and molecular dynamics. Antiviral Research 2017, 137, 151–164.

19. Liu, H., Okazaki, S., Shinoda, W. Heteroaryldihydropyrimidines alter capsid assembly by adjusting the binding affinity and pattern of the hepatitis B virus core protein. Journal of Chemical Information and Modeling 2019, 59, 5104–5110.

20. Pavlova, A., Bassit, L., Cox, B.D., Korablyov, M., Chipot, C., Verma, K., Russell, O.O., Schinazi, R.F., Gumbart, J.C. Mechanism of action of HBV capsid assembly modulators predicted from binding to early assembly intermediates. bioRxiv 2020, [https://www.biorxiv.org/content/early/2020/03 doi:10.1101/2020.03.23.002527.

21. Leaver-Fay, A., Tyka, M., Lewis, S.M., Lange, O.F., Thompson, J., Jacak, R., Kaufman, K.W., Renfrew, P.D., Smith, C.A., Sheffler, W., others. ROSETTA3: an object-oriented software suite for the simulation and design of macromolecules. In Methods in enzymology; Elsevier, 2011; Vol. 487, pp. 545–574.

22. Goh, B.C., Perilla, J.R., England, M.R., Heyrana, K.J., Craven, R.C., Schulten, K. Atomic modeling of an immature retroviral lattice using molecular dynamics and mutagenesis. Structure 2015, 23, 1414–1425.

23. Dolinsky, T.J., Czodrowski, P., Li, H., Nielsen, J.E., Jensen, J.H., Klebe, G., Baker, N.A. PDB2PQR: Expanding and upgrading automated preparation of biomolecular structures for molecular simulations. Nucleic Acids Research 2007, 35, W522–W525.

24. Humphrey, W., Dalke, A., Schulten, K., others. VMD: Visual Molecular Dynamics. Journal of Molecular Graphics 1996, 14, 33–38.

25. Jorgensen, W.L., Chandrasekhar, J., Madura, J.D., Impey, R.W., Klein, M.L. Comparison of simple potential functions for simulating liquid water. The Journal of Chemical Physics 1983, 79, 926–935.

26. Best, R.B., Zhu, X., Shim, J., Lopes, P.E., Mittal, J., Feig, M., MacKerell Jr, A.D. Optimization of the additive CHARMM all-atom protein force field targeting improved sampling of the backbone φ, ψ and side-chain χ1 and χ2 dihedral angles. Journal of Chemical Theory and Computation 2012, 8, 3257–3273.

27. Vanommeslaeghe, K., Hatcher, E., Acharya, C., Kundu, S., Zhong, S., Shim, J., Darian, E., Guvench, O., Lopes, P., Vorobyov, I., others. CHARMM general force field: A force field for drug-like molecules compatible with the CHARMM all-atom additive biological force fields. Journal of Computational Chemistry 2010, 31, 671–690.

28. Mayne, C.G., Saam, J., Schulten, K., Tajkhorshid, E., Gumbart, J.C. Rapid parameterization of small molecules using the force field toolkit. Journal of Computational Chemistry 2013, 34, 2757– 2770.

29. Frisch, M., Trucks, G., Schlegel, H., Scuseria, G., Robb, M., Cheeseman, J., Scalmani, G., Barone, V., Mennucci, B., Petersson, G., others. Gaussian09. Gaussian, Inc., Wallingford, CT 2009.

30. Pang, Y.T., Pavlova, A., Tajkhorshid, E., Gumbart, J.C. Parameterization of a drug molecule with a halogen σ-hole particle using ffTK: Implementation, testing, and comparison. The Journal of Chemical Physics 2020, 153, 164104.

31. Phillips, J.C., Braun, R., Wang, W., Gumbart, J., Tajkhorshid, E., Villa, E., Chipot, C., Skeel, R.D., Kale, L., Schulten, K. Scalable molecular dynamics with NAMD. Journal of Computational Chemistry 2005, 26, 1781–1802.

32. Bourne, C.R., Finn, M., Zlotnick, A. Global structural changes in hepatitis B virus capsids induced by the assembly effector HAP1. Journal of Virology 2006, 80, 11055–11061.

33. Frishman, D., Argos, P. Knowledge-based protein secondary structure assignment. Proteins: Structure, Function, and Bioinformatics 1995, 23, 566–579.

34. Perilla, J.R., Schulten, K. Physical properties of the HIV-1 capsid from all-atom molecular dynamics simulations. Nature Communications 2017, 8, 1–10.

35. Bakan, A., Meireles, L.M., Bahar, I. ProDy: Protein dynamics inferred from theory and experiments. Bioinformatics 2011, 27, 1575–1577.

36. Eargle, J., Luthey-Schulten, Z. NetworkView: 3D display and analysis of protein· RNA interaction networks. Bioinformatics 2012, 28, 3000–3001.

37. Dahl, A.C.E., Chavent, M., Sansom, M.S. Bendix: Intuitive helix geometry analysis and abstraction. Bioinformatics 2012, 28, 2193–2194.

38. Böttcher, B., Nassal, M. Structure of mutant hepatitis B core protein capsids with premature secretion phenotype. Journal of Molecular Biology 2018, 430, 4941–4954.

39. Feher, V.A., Durrant, J.D., Van Wart, A.T., Amaro, R.E. Computational approaches to mapping allosteric pathways. Current Opinion in Structural Biology 2014, 25, 98–103.

40. Zhao, Z., Wang, J.C.Y., Gonzalez-Gutierrez, G., Venkatakrishnan, B., Asor, R., Khaykelson, D., Raviv, U., Zlotnick, A. Structural differences between the woodchuck hepatitis virus core protein in the dimer and capsid states are consistent with entropic and conformational regulation of assembly. Journal of Virology 2019, 93.

41. Watts, N.R., Conway, J.F., Cheng, N., Stahl, S.J., Belnap, D.M., Steven, A.C., Wingfield, P.T. The morphogenic linker peptide of HBV capsid protein forms a mobile array on the interior surface. The EMBO Journal 2002, 21, 876–884.

42. Hilmer, J.K., Zlotnick, A., Bothner, B. Conformational equilibria and rates of localized motion within hepatitis B virus capsids. Journal of Molecular Biology 2008, 375, 581–594.

43. Venkatakrishnan, B., Katen, S.P., Francis, S., Chirapu, S., Finn, M., Zlotnick, A. Hepatitis B virus capsids have diverse structural responses to small-molecule ligands bound to the heteroaryldihydropyrimidine pocket. Journal of Virology 2016, 90, 3994–4004.

44. Patterson, A., Zhao, Z., Waymire, E., Zlotnick, A., Bothner, B. Dynamics of Hepatitis B virus capsid protein dimer regulate assembly through an allosteric network. ACS Chemical Biology 2020, 15, 2273–2280.

45. Bereszczak, J.Z., Watts, N.R., Wingfield, P.T., Steven, A.C., Heck, A.J. Assessment of differences in the conformational flexibility of hepatitis B virus core-antigen and e-antigen by hydrogen deuterium exchange-mass spectrometry. Protein Science 2014, 23, 884–896.

46. Altschul, S.F., Gish, W., Miller, W., Myers, E.W., Lipman, D.J. Basic local alignment search tool. Journal of Molecular Biology 1990, 215, 403–410.

47. Selzer, L., Katen, S.P., Zlotnick, A. The hepatitis B virus core protein intradimer interface modulates capsid assembly and stability. Biochemistry 2014, 53, 5496–5504.

48. Ceres, P., Stray, S.J., Zlotnick, A. Hepatitis B virus capsid assembly is enhanced by naturally occurring mutation F97L. Journal of Virology 2004, 78, 9538–9543.

